# The Microtubule Associated Protein Tau Regulates KIF1A Pausing Behavior and Motility

**DOI:** 10.1101/2021.08.11.455914

**Authors:** DV Lessard, CL Berger

**Affiliations:** University of Vermont

**Keywords:** KIF1A, Tau, C-terminal tail, kinesin, microtubule, axonal transport, neurodegenerative disease, microtubule associated protein (MAP)

## Abstract

Many neurodegenerative diseases result from dysfunction of axonal transport, a highly regulated cellular process responsible for site-specific neuronal cargo delivery. The kinesin-3 family member KIF1A is a key mediator of this process by facilitating long-distance cargo delivery in a spatiotemporally regulated manner. While misregulation of KIF1A cargo delivery is observed in many neurodegenerative diseases, the regulatory mechanisms responsible for KIF1A cargo transport are largely unexplored. Our lab has recently characterized a mechanism for a unique pausing behavior of KIF1A in between processive segments on the microtubule. This behavior, mediated through an interaction between the KIF1A K-loop and the polyglutamylated C-terminal tails of tubulin, helps us further understand how KIF1A conducts long-range cargo transport. However, how this pausing behavior is influenced by other regulatory factors on the microtubule is an unexplored concept. The microtubule associated protein Tau is one potential regulator, as altered Tau function is a pathological marker in many neurodegenerative diseases. However, while the effect of Tau on kinesin-1 and -2 has been extensively characterized, its role in regulating KIF1A transport is greatly unexplored at the behavioral level. Using single-molecule imaging, we have identified Tau-mediated regulation of KIF1A pausing behavior and motility. Specifically, our findings imply a competitive interaction between Tau and KIF1A for the C-terminal tails of tubulin. We introduce a new mechanism of Tau-mediated kinesin regulation by inhibiting the ability of KIF1A to use C-terminal tail reliant pauses to connect multiple processive segments into a longer run length. Moreover, we have correlated this regulatory mechanism to the behavioral dynamics of Tau, further elucidating the function of Tau diffusive and static behavioral state on the microtubule surface. In summary, we introduce a new mechanism of Tau-mediated motility regulation, providing insight on how disruptions in axonal transport can lead to disease state pathology.

**SIGNIFICANCE:** KIF1A mediated cargo transport is essential in many cellular processes such as axonal transport and neuronal development. Defects in KIF1A transport have been implicated in neurodegenerative diseases including Alzheimer’s disease and frontotemporal dementia. However, the mechanism of KIF1A’s pathological misregulation remains elusive, highlighting the importance of identifying regulators of KIF1A function. The microtubule associated protein Tau is an attractive potential regulator of KIF1A motility as Tau dysfunction is a hallmark of these neurodegenerative diseases. Here, we demonstrate a direct connection between Tau and KIF1A motility, revealing a unique form of Tau-mediated regulation of axonal transport. Our results provide a molecular foundation for understanding the role of motor protein misregulation in neurodegenerative disease progression.

## Introduction

Impaired cellular cargo trafficking is a hallmark of many neurodegenerative diseases, such as Alzheimer’s disease (AD) and frontotemporal dementia (FTD) [1-4]. This disease-state pathology implicates a deficiency in axonal transport, a critical process for neuronal viability and function, involving the spatiotemporal trafficking of cellular cargo along the length of the axon. A key mediator of this process is the kinesin-3 family member KIF1A, known to transport cargo that must travel great magnitudes of distance down the axon, such as dense core vesicles [5] and synaptic vesicle proteins [6, 7]. Recent discoveries uncovering the motor specific behavior of KIF1A, such as its “superprocessive” motility [8, 9] and characteristic pausing in-between processive segments of motility [10], have provided significant insight as to how this motor is able to transport cargo such extreme magnitudes of distance in comparison to other kinesin motors. Moreover, KIF1A’s motor specific behavior and motility must be tightly regulated to achieve efficient spatiotemporal cargo delivery within the neuron. The systemic importance of proper KIF1A regulation is demonstrated in neurodegenerative diseases presenting with KIF1A cargo mislocalization, such as AD and FTD [11-13]. However, the mechanisms contributing towards pathological KIF1A cargo delivery are greatly unexplored, stemming from our lack of knowledge of mechanisms that regulate KIF1A motility.

The complex landscape of axonal microtubules may provide a regulatory mechanism for the spatiotemporal delivery of KIF1A cargo, via the presence of microtubule associated proteins (MAPs) that bind to the microtubule surface. Specifically, the neuronal MAP Tau is an attractive potential regulator of KIF1A, as Tau has been shown to differentially regulate the motility of specific kinesin families. Initial observations of the highly characterized kinesin-1 family of motors revealed that, upon encountering microtubule bound Tau, kinesin-1 motors prematurely dissociate from the microtubule surface [14, 15]. Our lab has further explored Tau’s inhibition of kinesin-1 motility, discovering that a pathologically relevant phosphomimetic mutation of Tau leads to an intermediary level of kinesin-1 motility inhibition, as well as showing that Tau’s inhibition of kinesin-1 depends on the nucleotide binding state of microtubules [16, 17]). Contrastingly, our lab has also shown that unlike kinesin-1, kinesin-2 motility is insensitive to Tau [18, 19]. Despite the complex array of Tau’s regulatory mechanisms on other kinesin family members, few studies have explored the effects of Tau on dimeric KIF1A motility parameters. Preexisting studies have advanced our understanding of Tau’s ability to alter KIF1A landing rate and processive characteristics on microtubules [20, 21]. Because of these initial findings, it is imperative that the relationship between KIF1A and Tau be more thoroughly characterized at a molecular level to build the cellular framework needed to understand Tau’s effect on KIF1A cargo transport in a neurodegenerative setting. Pathologically, irregular Tau function in AD and FTD further emphasizes the importance of defining this relationship [3, 4, 22-24], insinuating that Tau’s loss-of-function could sacrifice a critical regulatory mechanism of KIF1A cargo delivery. However, the lack of research on this topic highlights a gap in knowledge halting our understanding of neurodegenerative disease progression.

Intriguingly, both KIF1A and Tau demonstrate unique behavioral interactions with the microtubule surface. Tau binds to the microtubule surface both statically and diffusively in an equilibrium that is isoform specific [25, 26]. It has been further shown that Tau relies on the tubulin C-terminal tails (CTTs) to facilitate specifically diffusive binding behavior [25]. Based on this interaction, the isoform-dependent equilibrium of Tau’s static and diffusive binding is an important parameter to consider when assessing Tau’s regulation of kinesin motors. For example, our lab has shown that kinesin-1 motors are more severely inhibited by the static binding state of Tau, not the diffusive binding state of Tau [16, 18, 19]. Yet, how Tau’s isoform-specific binding equilibrium may regulate KIF1A is a concept entirely unexplored.

Much like Tau, the CTTs of tubulin are an essential structure for KIF1A function. The KIF1A/CTT interaction is largely mediated by an electrostatic “tethering” between KIF1A’s K-loop, a positively charged surface loop in the motor domain, and post-translational addition of negatively charged glutamic acid residues to the CTTs, a signature of neuronal tubulin [10]. This K-loop/CTT interaction is known to be important for many KIF1A characteristics such as landing rate and facilitating diffusive events along the microtubule surface in the weakly-bound ADP state [9, 27]. In addition to these findings, our lab has recently described the characteristic pausing behavior of KIF1A, revealing that KIF1A relies on the CTTs to engage in pauses in-between segments of processive movement on the microtubule [10]. Considering that both KIF1A and diffusively bound Tau interact with the microtubule C-terminal tails, we hypothesized that diffusive Tau has the potential to act as regulator of KIF1A motility by interacting with the CTT during KIF1A pausing (Figure 1). Using single-molecule total internal reflection fluorescence microscopy, we tested our hypothesis by quantifying the motility and behavioral response of KIF1A on Tau decorated microtubules. Our findings uncovered a new mechanism of Tau-mediated kinesin motor inhibition. From this, we present the discovery of diffusive Tau’s inhibition of KIF1A motility during KIF1A pausing.

**Figure 1.**
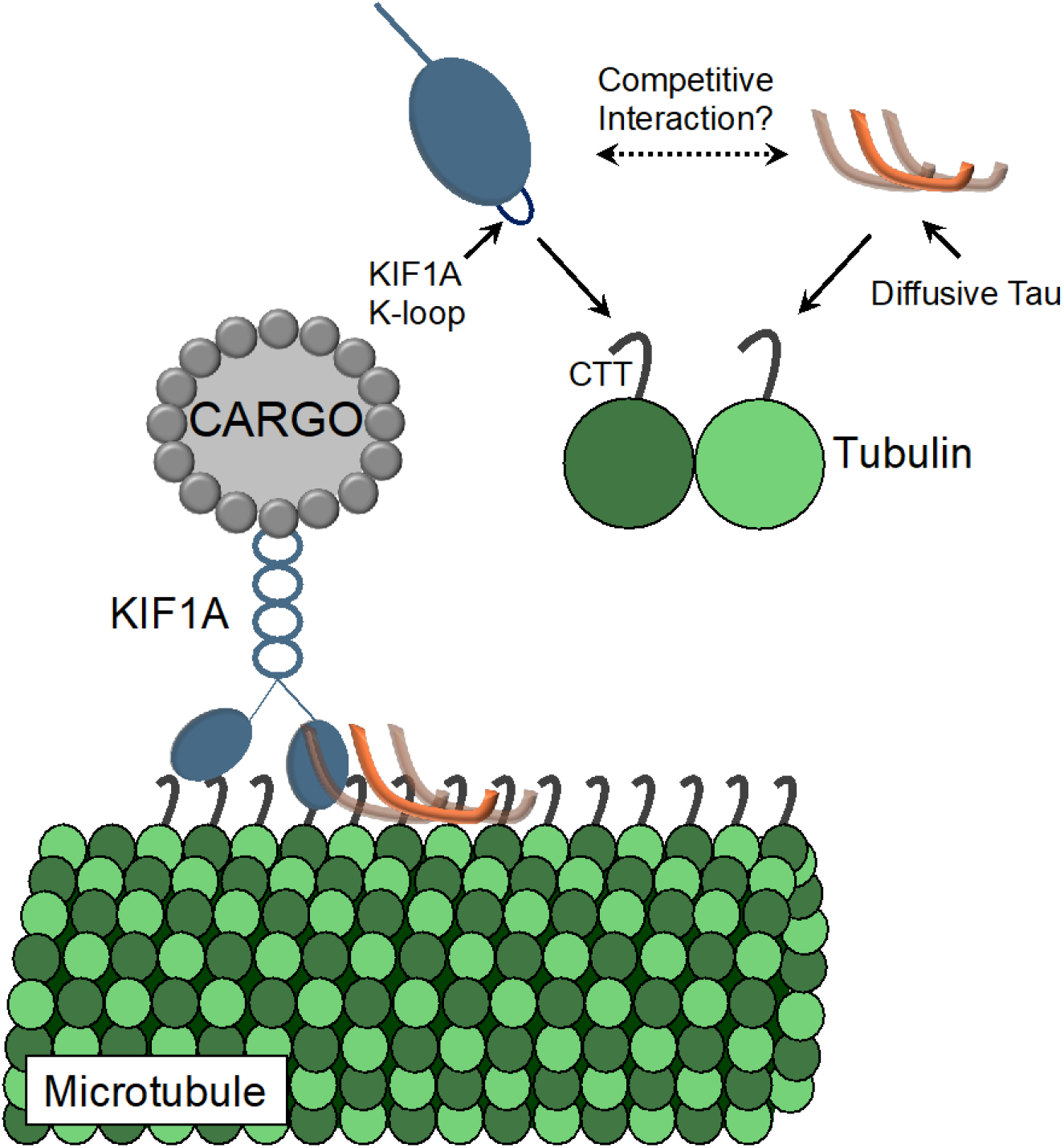
Graphical representation of experimental hypothesis. Previous literature has detailed that both KIF1A and Tau interact with the C-terminal tails (CTT) of tubulin subunits. The interaction between KIF1A’s “K-loop” and the polyglutamylated CTTs has been shown to facilitate KIF1A’s recently characterized pausing behavior [10]. Each Tau isoform has a characteristic behavioral equilibrium between a static and a diffusing binding state. The diffusive binding state of Tau is achieved through an interaction with the CTTs [25]. As both proteins rely on interactions with the CTTs to exhibit protein-specific behavior, we hypothesis that the diffusive binding state of Tau engaged with the CTTs inhibits KIF1A pausing.

## MATERIALS AND METHODS

### Microtubule preparation and labelling

Tubulin was isolated from bovine brains donated from Vermont Livestock Slaughter & Processing (Ferrisburgh, VT) as previous described [10]. Isolated tubulin was clarified using an Optima TLX Ultracentrifuge (Beckman, Pasadena, CA) at 95,000 rpm for 20 minutes at 4°C. Clarified tubulin was read at 280nm in a spectrophotomer and the extinction coefficient of 115,000 cm^-1^ M^-1^ was used to determine concentration. Clarified tubulin was supplemented with 1mM GTP (Sigma-Aldrich, St. Luois, MO) and labelled, stabilized, and diluted as previously described [10]. For microtubule C-terminal tail removal, microtubules were treated with Subtilisin A (Sigma Aldrich) as previously described [10]. For experiments containing Tau, tubulin polymerization was performed as described above, in the absence of labeled tubulin. Polymerized microtubules were incubated with 200 nM, 100 nM or 50 nM Alexa 647 labeled Tau (1:5, 1:10 or 1:20 Tau to tubulin ratio) for an additional 20 minutes at 37°C. Tau and microtubules mixture was centrifuged at room temperature for 30 minutes at 15,000 rpm. The pellet was then resuspended in Motility Buffer. To ensure the same amount of 3RS-Tau decoration on untreated vs. subtilisin-treated microtubules, 3RS-Tau’s fraction bound at our experimental microtubule concentration was determined as previously described [16] and adjusted accordingly. Adjusted concentrations are noted in figure legends when applicable.

### Protein expression and labelling

Wild-type (WT) 3RS-and 4RL-Tau constructs were expressed in BL21 CodonPlus(DE3)-RP Escherichia coli cells (Stratagene, La Jolla, CA) and purified as previously described [17, 26]. Purified protein was dialyzed in 1x BRB80, and concentration was determined using the Bicinchoninic Acid (BCA) Protein Assay (Pierce, Rockford, IL) using WT 3RS-Tau as a standard. To label Tau, protein was incubated with a 10-fold molar excess of dithiothreitol (DTT) at room temperature for 2 hours. DTT was then removed using a 2 ml 7K MWCO Zeba spin desalting column (Pierce, Rockford, IL). Protein was then incubated with fivefold molar excess of Alexa 647 C5 Maleimide (Invitrogen Molecular Probes, Carlsbad, CA) overnight at room temperature. Excess fluor was removed using the previously mentioned Zeba desalting columns. To determine the labeling efficiency, Alexa Fluor 647 concentration was determined using an extinction coefficient of 270,000 cm^-1^ M^-1^ at 495 nm using a 640 Spectrophotometer. Labeled protein concentration was determined using a BCA assay. KIF1A(1-393)-LZ-3xmCitrine motors (a generous gift of Dr. Kristen Verhey, University of Michigan, Ann Arbor, MI) were expressed in COS-7 monkey kidney fibroblasts (American Type Culture Collection, Mansses, VA) as previously described [10].

### In vitro single-molecule TIRF

Flow chambers used in *in vitro* TIRF experiments were constructed as previously described [16]. Flow chambers were incubated with monoclonal anti-β III (neuronal) antibodies at 33 μg/ml for 5 minutes, then washed twice with 0.5 mg/ml bovine serum albumin (BSA; Sigma Aldrich) and incubated for 2 minutes. 1 μM of microtubules (all experimental conditions) were administered and incubated for 8 minutes. Non-adherent microtubules were removed with a MB wash supplemented with 20 μM paclitaxel. Kinesin motors in Motility Buffer (MB; 12 mM PIPES, 1 mM MgCl_2_, 1 mM EGTA, supplemented with 20 μM paclitaxel, 10 mM DTT, 1 mM MgCl_2_, 10 mg/ml BSA, 2 mM ATP and an oxygen scavenger system [5.8 mg/ml glucose, 0.045 mg/ml catalase, and 0.067 mg/ml glucose oxidase; Sigma Aldrich]), supplemented with 2 mM ATP, were added to the flow cell just before image acquisition. For landing rate assays, *in vitro* KIF1A motility assays were prepared as described above, with the only change being that MB was supplemented with 2 mM ADP. As kinesin-1 motility has been extensively characterized in *in vitro* reconstituted motility experiments in our lab and many others, we use this construct on untreated, stabilized microtubules as an internal control to ensure that our experimental system is functioning optimally. Control Drosophila melanogaster biotin-tagged kinesin-1 motors were labeled with streptavidin-conjugated Qdot 655 (Life Technologies, Carlsbad, CA) at a 1:4 motor:Qdot ratio as previously described [16, 19].

### In vitro fluorescence based Tau binding assay

To assess the affinity of Tau for subtilisin treated microtubules, we performed a TIRF binding assay, as previously described [16]. In brief, unlabeled subtilisin-treated microtubules were diluted to a working concentration of 1.5 μM, adhered to flow chambers as previously described, and washed with MB to remove non-adherent microtubules. Next, 10 nM of Alexa 647 labeled Tau was flowed in and imaged at 10 frames/s for 20 frames. This process was repeated with increasing concentrations of Alexa 647 labeled Tau and conducted on the previously described TIRF set up.

### Data Analysis

#### In vitro motility assays

Motility events were analyzed as previously reported [18, 19]. In brief, overall run length motility data was measured using the ImageJ (v. 2.0.0, National Institute of Health, Bethesda, MD) MTrackJ plug-in, for a frame-by-frame quantification of KIF1A motility. KIF1A’s overall run length is defined as the summation of continuous run lengths and pausing events during a single motility event on the microtubule. Continuous run length is defined as segments where, within a single event, KIF1A is moving at a constant velocity. Pauses are defined as segments in where the average velocity is less than 0.2 μm/s over three frames or more. Processive events were identified at the boundaries of pausing events [10]. Kymographs of motor motility were created using the MultipleKymograph ImageJ plugin, with a set line thickness of 3. To correct for microtubule track length effects on motor motility, overall/processive run length data was resampled to generate cumulative frequency plots (99% CI), using previously reported methods [28]. Statistical significance between corrected motor run length and speed data sets was determined using a paired t-test. To determine the landing rate of KIF1A on various microtubule conditions, the total number of motor landing events on the microtubule was divided by the length of the microtubule and further divided by the duration of the movie (events/μm/minute).

#### Tau Binding Assay

Differences in 3RS-Tau binding affinity on untreated vs. subtilisin-treated microtubules were determined using a TIRF Tau binding assays were analyzed as previously described [16]. In summary, using the MultiMeasure tool in FIJI, the average intensity of Tau per unit length of microtubule (Avg I) was measured for each frame. Average intensity was normalized to B_max_ for each respective concentration, then plotted against [Tau] to generate a binding curve and fit to one-site-specific binding with Hill slope. From this fit, a Hill coefficient (h) and *K*_*D*_ were determined. From this, the fraction of 3RS-Tau on subtilisin-treated microtubules was determined as previously described [16].

## RESULTS

### 3RS-Tau inhibits KIF1A overall run length, but not continuous run length

Using TIRF microscopy, we first assessed the inhibitory effect of 3RS-Tau, a well characterized isoform in terms of kinesin motility regulation that favors the static binding state (Figure 2A) [14-16, 18, 19, 26], on single-molecule KIF1A motility. KIF1A’s motility was first investigated on paclitaxel-stabilized microtubules in the presence of Alexa 647 labeled wild-type (WT) 3RS-Tau at three concentrations (50 nM, 100 nM, or 200 nM 3RS-Tau ; 1 μM microtubules). In the absence of Tau, KIF1A’s overall run length was 6.84 ± 2.07 μm (Figure 2C, Table 1) and overall speed was 1.37 ± 0.42 μm/s (Figure S1, Table 1), supporting past reports of this motor’s superprocessivity and consistent with our previous work characterizing KIF1A’s pausing behavior [8, 10]. In the presence of 3RS-Tau KIF1A’s overall run length significantly decreased in a dosage dependent manner (50 nM 3RS-Tau; 4.65 ± 1.83 μm, 100 nM 3RS-Tau; 3.84 ± 1.50 μm, 200 nM 3RS-Tau; 3.31 ± 1.06 μm) (Figure 2C, Figure S3, Table 1), while overall speed increased with the addition of Tau (50 nM 3RS-Tau; 1.38 ± 0.34 μm/s, 100 nM 3RS-Tau; 1.40 ± 0.37 μm/s, 200 nM 3RS-Tau; 1.59 ± 0.39 μm/s) (Figure S1, Table 1). These results provide direct evidence of 3RS-Tau’s ability to inhibit KIF1A motility by both decreasing the overall run length while simultaneously increasing overall speed.

**Figure 2.**
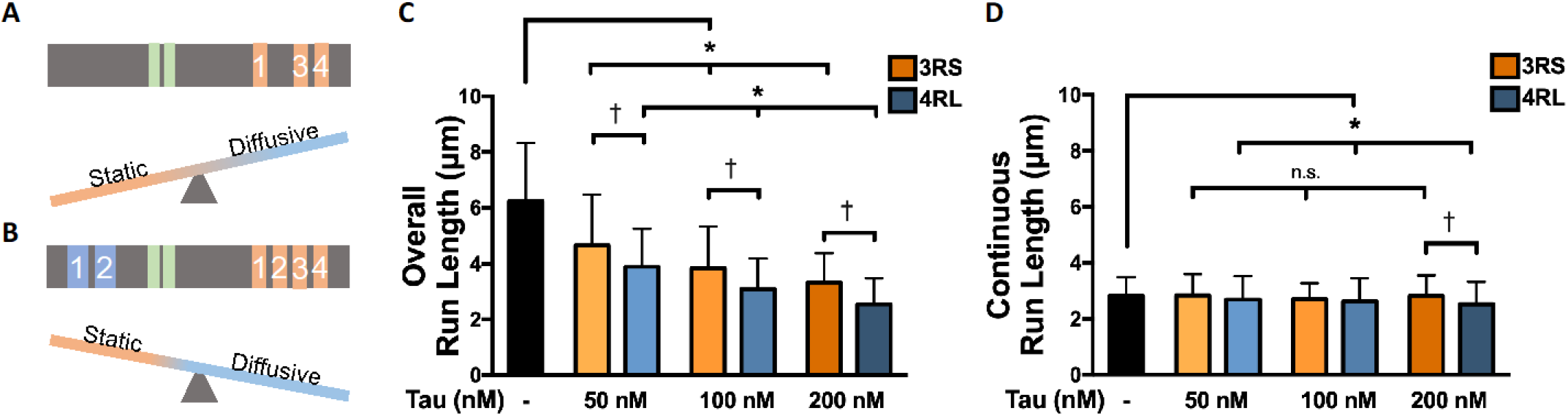
4RL-Tau more strongly reduces KIF1A run length than 3RS-Tau. **A)** Cartoon depicting structural and behavioral characteristics of 3RS-Tau. Top: Linearized 3RS-Tau protein structure, featuring a proline rich region (green) and three microtubule binding repeats (orange). Bottom: The behavioral equilibrium of 3RS-Tau favors the static binding state. **B)** Cartoon depicting structural and behavioral characteristics of 4RL-Tau. Top: Linearized 4RL-Tau protein structure, featuring two acidic inserts (blue) a proline rich region (green) and four microtubule binding repeats (orange). Bottom: The behavioral equilibrium of 4RL-Tau favors the diffusive binding state. **C)** The overall run length (ORL) of KIF1A was reduced upon the addition of either 3RS-or 4RL-Tau in a dosage dependent manner. When 3RS-and 4RL-Tau are compared at specific concentrations, it is revealed that 4RL is more inhibitory than 3RS-Tau. **D)** Continuous run length was not significantly reduced between 3RS-and 4RL-Tau at 50 nM and 100 nM concentrations, but was significantly reduced at the highest Tau concentration (200 nM). Run length values are reported as mean ± standard deviation and were calculated as previously reported [28]. *^,ƚ^ p <0.05

**Table 1.**
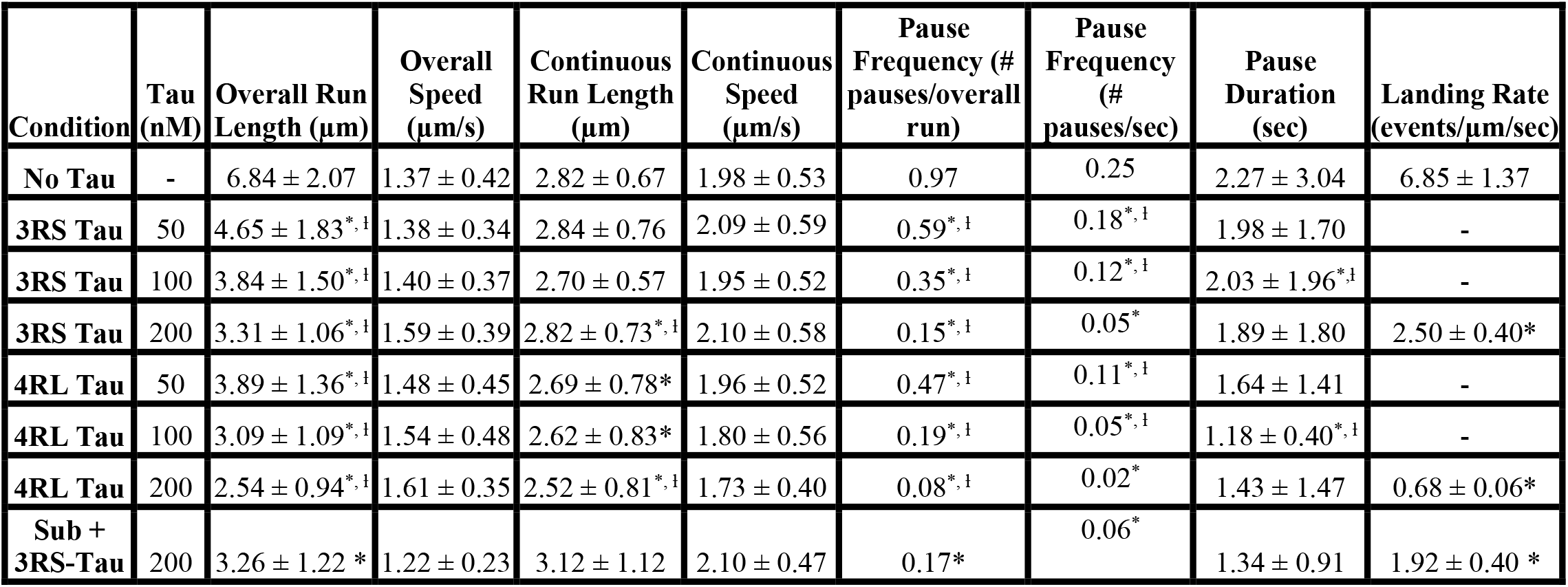
Summary of KIF1A motility and behavior on microtubules (1 μM) across all conditions. Metrics are reported as Mean ± SD. * = p<0.05 relative to KIF1A on “No Tau” MTs. ƚ = p<0.05 relative to 3RS-vs. 4RL-Tau at each respective concentration.

| Condition | Tau (nM) | Overall Run Length (μm) | Overall Speed (μm/s) | Continuous Run Length (μm) | Continuous Speed (μm/s) | Pause Frequency (# pauses/overall run) | Pause Frequency (# pauses/sec) | Pause Duration (sec) | Landing Rate (events/μm/sec) |
| --- | --- | --- | --- | --- | --- | --- | --- | --- | --- |
| No Tau | - | 6.84 ± 2.07 | 1.37 ± 0.42 | 2.82 ± 0.67 | 1.98 ± 0.53 | 0.97 | 0.25 | 2.27 ± 3.04 | 6.85 ± 1.37 |
| 3RS Tau | 50 | 4.65 ± 1.83<sup>*,1</sup> | 1.38 ± 0.34 | 2.84 ± 0.76 | 2.09 ± 0.59 | 0.59<sup>*,1</sup> | 0.18<sup>*,1</sup> | 1.98 ± 1.70 | - |
| 3RS Tau | 100 | 3.84 ± 1.50<sup>*,1</sup> | 1.40 ± 0.37 | 2.70 ± 0.57 | 1.95 ± 0.52 | 0.35<sup>*,1</sup> | 0.12<sup>*,1</sup> | 2.03 ± 1.96<sup>*,1</sup> | - |
| 3RS Tau | 200 | 3.31 ± 1.06<sup>*,1</sup> | 1.59 ± 0.39 | 2.82 ± 0.73<sup>*,1</sup> | 2.10 ± 0.58 | 0.15<sup>*,1</sup> | 0.05<sup>*</sup> | 1.89 ± 1.80 | 2.50 ± 0.40<sup>*</sup> |
| 4RL Tau | 50 | 3.89 ± 1.36<sup>*,1</sup> | 1.48 ± 0.45 | 2.69 ± 0.78<sup>*</sup> | 1.96 ± 0.52 | 0.47<sup>*,1</sup> | 0.11<sup>*,1</sup> | 1.64 ± 1.41 | - |
| 4RL Tau | 100 | 3.09 ± 1.09<sup>*,1</sup> | 1.54 ± 0.48 | 2.62 ± 0.83<sup>*</sup> | 1.80 ± 0.56 | 0.19<sup>*,1</sup> | 0.05<sup>*,1</sup> | 1.18 ± 0.40<sup>*,1</sup> | - |
| 4RL Tau | 200 | 2.54 ± 0.94<sup>*,1</sup> | 1.61 ± 0.35 | 2.52 ± 0.81<sup>*,1</sup> | 1.73 ± 0.40 | 0.08<sup>*,1</sup> | 0.02<sup>*</sup> | 1.43 ± 1.47 | 0.68 ± 0.06<sup>*</sup> |
| Sub + 3RS-Tau | 200 | 3.26 ± 1.22 <sup>*</sup> | 1.22 ± 0.23 | 3.12 ± 1.12 | 2.10 ± 0.47 | 0.17<sup>*</sup> | 0.06<sup>*</sup> | 1.34 ± 0.91 | 1.92 ± 0.40 <sup>*</sup> |

To further investigate the process by which *overall* run length is reduced by 3RS-Tau, we next assessed changes in KIF1A’s *continuous* run length, a main component of the overall run length measurement [10]. In contrast to the trends observed with overall run length, there was no significant reduction of KIF1A’s continuous run length between our control microtubules (no Tau), or at any concentration of 3RS-Tau (No Tau; 2.82 ± 0.73 μm, 50 nM 3RS-Tau; 2.84 ± 0.76 μm, 100 nM 3RS-Tau; 2.70 ± 0.57 μm, 200 nM 3RS-Tau; 2.82 ± 0.73 μm) (Figure 2D, Figure S4, Table 1). Following suit, there was also no change in continuous speed at any 3RS-Tau concentration (50 nM 3RS-Tau; 2.09 ± 0.59 μm/s, 100 nM 3RS-Tau; 1.95 ± 0.52 μm/s, 200 nM 3RS-Tau; 2.10 ± 0.58 μm/s) when compared to our control (Figure S2, Table 1). From these experiments we can conclude that, while 3RS-Tau inhibits KIF1A’s overall run length and speed, it does not do so by inhibiting KIF1A’s continuous run length or speed.

### 4RL-Tau is more inhibitory than 3RS-Tau on KIF1A motilit

To compare the KIF1A inhibition via two Tau isoforms with contrasting binding equilibriums, we next turned to Alexa 647 labeled WT 4RL-Tau, an isoform that favors the diffusive binding state (Figure 2A-B). Similar to 3RS-Tau, we observed a dosage dependent decrease of KIF1A overall run length (50 nM 4RL-Tau; 3.89 ± 1.36 μm, 100 nM 4RL-Tau; 3.09 ± 1.09 μm, 200 nM 4RL-Tau; 2.54 ± 0.94 μm) (Figure 2C, Figure S3, Table 1) and a dosage dependent increase in overall speed (50 nM 4RL-Tau; 1.48 ± 0.45 μm/s, 100 nM 4RL-Tau; 1.54 ± 0.48 μm/s, 200 nM 4RL-Tau; 1.61 ± 0.35 μm/s) in the presence of 4RL-Tau. However, when KIF1A’s overall run length was compared between 3RS-and 4RL-Tau, KIF1A’s overall run length was significantly shorter on 4RL-Tau coated microtubules every Tau concentration (Figure 2C, Figure S3, Table 1). The culmination of these results revealed to us that 4RL-Tau is statistically more inhibitory than 3RS-Tau on KIF1A overall run length across a dosage of Tau concentrations.

To further investigate how 4RL-Tau is more inhibitory on KIF1A overall run length, we compared the effects of 3RS-vs 4RL-Tau on KIF1A continuous run length. Like 3RS-Tau, there was no significant difference between KIF1A continuous run length at any concentration of 4RL-Tau (50 nM 4RL-Tau; 2.69 ± 0.78 μm, 100 nM 4RL-Tau; 2.62 ± 0.83 μm, 200 nM 4RL-Tau; 2.52 ± 0.81μm) (Figure 2D, Figure S4, Table 1) however all 4RL-Tau concentrations significantly reduced KIF1A continuous run length when compared to control microtubules. Unlike 3RS-Tau, we observed a dosage dependent decrease in continuous speed (50 nM 4RL-Tau; 1.96 ± 0.52 μm/s, 100 nM 4RL-Tau; 1.80 ± 0.56 μm/s, 200 nM 4RL-Tau; 1.73 ± 0.40 μm/s) when compared to our control (Figure S2, Table 1). Additionally, at our highest Tau concentration (200 nM), KIF1A continuous run length is significantly shorter on 4RL-Tau coated microtubules than 3RS-Tau coated microtubules (Figure 2D, Figure S4, Table 1). Taken together, these results show that 4RL-Tau does not shorten KIF1A continuous run length between any of our experimental concentrations of 4RL-Tau. However, the changes in continuous speed as well as significant differences in continuous run length when compared to our control or 3RS-Tau at our highest experimental concentration support the idea that 4RL-Tau is more inhibitory than 3RS-Tau on KIF1A motility.

### KIF1A pausing behavior is inhibited by 3RS-and 4RL-Tau in a dosage dependent manner, and is more strongly inhibited by 4RL-Tau than 3RS-Tau

As referenced earlier, KIF1A’s overall run length is defined as the summation of continuous run lengths and pausing events during a single motility event on the microtubule [10]. With the effect of 3RS-and 4RL-Tau on KIF1A overall and continuous run length established, we next investigated Tau’s effect on KIF1A’s characteristic pausing behavior. KIF1A’s pausing behavior, and other quantifiable behaviors such as landing rate, are made possible through an interaction between the C-terminal tail structure of tubulin and the KIF1A K-loop [9, 10]. Moreover, Tau also relies on the C-terminal tails to engage in diffusive binding behavior [25]. The potential occupancy of both KIF1A and Tau on the tubulin C-terminal tails, combined with the observed inhibitory effect of 3RS-and 4RL-Tau on KIF1A’s overall run length, but not continuous run length, lead us to investigate KIF1A pausing as the point of Tau-mediated inhibition.

Upon addition of either 3RS-or 4RL-Tau, KIF1A pause frequency (# of pauses/overall run length) was drastically reduced (Figure 3A-B, Figure S6). Similar to what we observed regarding the influence of Tau on KIF1A’s overall run length, we observed a dosage dependent decrease on KIF1A pause frequency on both 3RS-(50 nM 3RS-Tau; 0.59 ± 0.06 pauses/ORL, 100 nM 3RS-Tau; 0.35 ± 0.07 pauses/ORL, 200 nM 3RS-Tau; 0.15 ± 0.03 pauses/ORL) and 4RL-Tau (50 nM 4RL -Tau; 0.47 ± 0.05 pauses/ORL, 100 nM 4RL -Tau; 0.19 ± 0.03 pauses/ORL, 200 nM 4RL -Tau; 0.08 ± 0.02 pauses/ORL) coated microtubules when compared to control microtubules (0.97 ± 0.07 pauses/ORL) (Figure 3C, Figure S5, Figure S6, Table 1). A reduction in KIF1A pause duration was observed both with 4RL-Tau at 100 nM concentration between 3RS-Tau (3RS; 2.03 ± 1.96 s, 4RL; 1.18 ± 0.40 s) as well as between 4RL-Tau at 100 nM concentration and our control microtubules (2.27 ± 3.04 s) (Figure 3D). However, it is important to consider that the extensive variability between conditions due to small sample sizes (less measurable pauses as Tau concentration increases) and large standard deviations is likely to skew this result.

**Figure 3.**
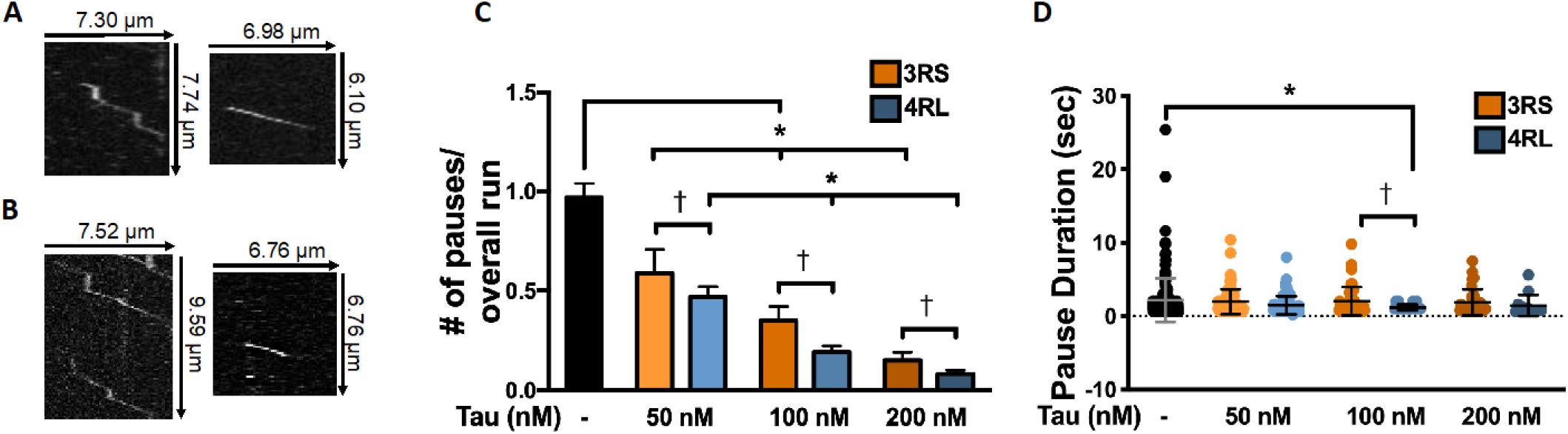
4RL-Tau more strongly inhibits KIF1A pausing behavior when compared to 3RS-Tau. **A)** Representative kymographs of KIF1A behavior on microtubules with no Tau (left) and 3RS-Tau (200 nM; right). **B)** Representative kymographs of KIF1A behavior on microtubules with no Tau (left) and 4RL-Tau (200 nM; right). **C**) KIF1A pause frequency (# of pauses/ overall run) decreases in a dosage-dependent manner upon addition of 3RS-and 4RL-Tau. At individual concentrations of Tau (50 nM, 100 nM, or 200 nM), 4RL-Tau significantly reduces the pause frequency of KIF1A when compared to 3RS-Tau. **D)** KIF1A pause duration did not significantly change upon addition of 3RS-or 4RL-Tau (with the exception of 100 nM 4RL-Tau). *^,ƚ^ p <0.05

In comparing kymographs of KIF1A motility we also observed a visually appreciable reduction in motor landing events on Tau-coated microtubules when compared to control microtubules, consistent with past literature that has highlighted the inhibitory nature of high-density Tau patches on KIF1A landing [20]. This, combined with KIF1A’s previous established reliance of the tubulin C-terminal tail for landing [9], lead us to explore potential differences in KIF1A landing rate in the absence and presence of 3RS-and 4RL-Tau. At our highest Tau concentration (200 nM), we observed a significant reduction in KIF1A landing rate on microtubules coated in either 3RS-(2.50 ± 0.40 events/μm/min) or 4RL-Tau (0.68 ± 0.06 events/μm/min) when compared to our control microtubules (6.35 ± 1.37 events/μm/min) (Figure 4).

**Figure 4.**
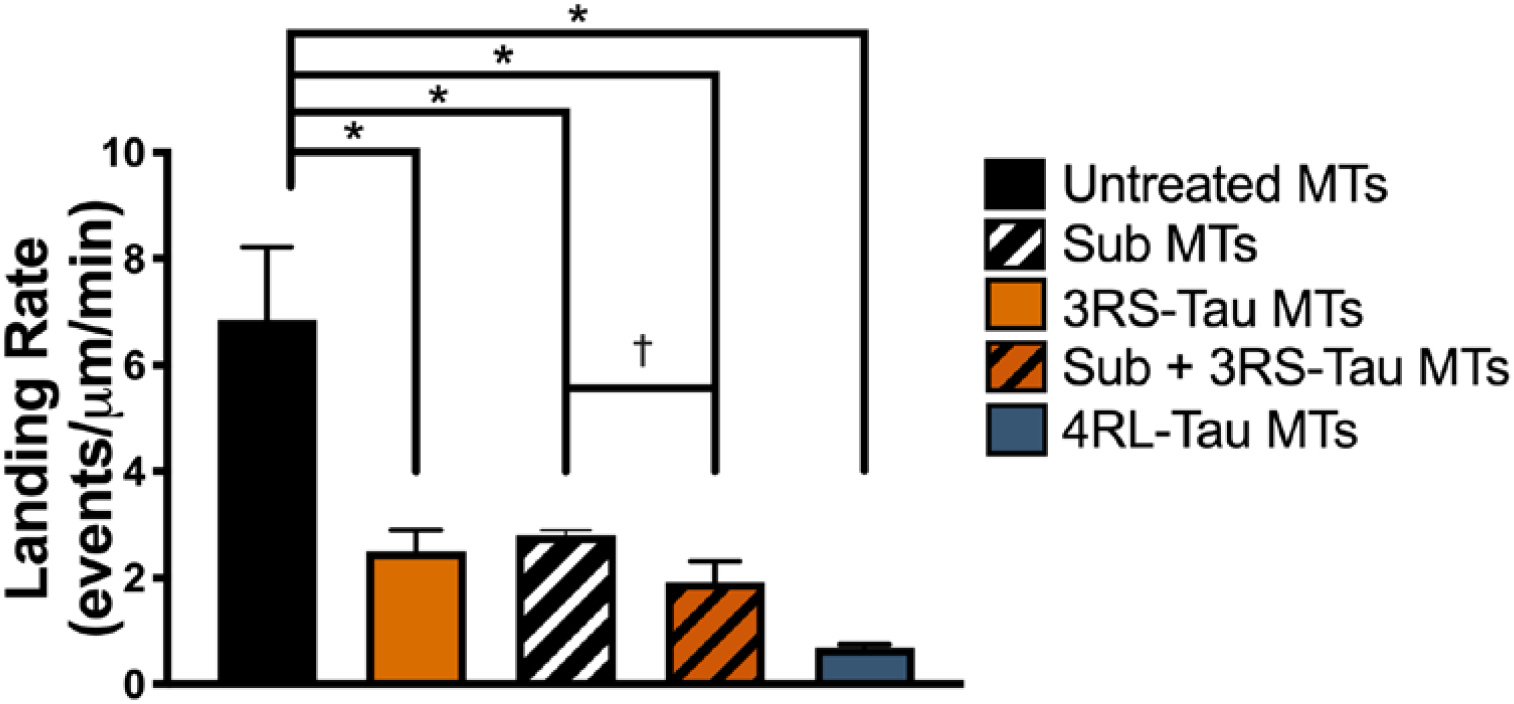
Quantification of KIF1A landing rate in the ADP state on indicated microtubule subsets. On undecorated, untreated microtubules, KIF1A had a landing rate of 6.85 ± 1.37 events/μm/min (N=971). Landing rate was significantly reduced with the addition of 3RS-Tau (2.50 ± 0.40 events/μm/min, N=846, 200 nM Tau), subtilisin treatment with the addition of 3RS-Tau (1.92 ± 0.40 events/μm/min, N=742, 200 nM Tau), or with the addition of 4RL-Tau (0.68 ± 0.06 events/μm/min, N=575, 200 nM Tau). Control subtilisin-treated microtubules (2.80 ± 0.42 events/μm/min, N=717) has been previously reported in Lessard *et al*., 2019 [10]* = p<0.001 relative to KIF1A on undecorated, untreated microtubules. ƚ = p<0.001 relative to KIF1A on undecorated, subtilisin-treated microtubules.

If KIF1A’s pausing is the point of Tau-mediated inhibition, then how might this relate to the characteristic binding equilibrium of each Tau isoform? Considering that Tau must also engage with the tubulin C-terminal tails to bind diffusively, and that 4RL-Tau favors the diffusive binding state, we further hypothesized that 4RL-Tau would be more inhibitory to KIF1A pausing than 3RS-Tau. This was confirmed when KIF1A’s pause frequency was compared between 3RS-and 4RL-Tau, revealing that KIF1A’s pause frequency was significantly reduced on 4RL-Tau coated microtubules at every Tau concentration (Figure 3C, Figure S5, Figure S6, Table 1). The culmination of these results support our idea that KIF1A pauses are heavily subjected to Tau-mediated inhibition. Furthermore, these findings insinuate that the diffusive binding state, not the static binding state, of Tau is inhibiting KIF1A pausing and is reducing the motors ability to resume a processive run after a pause, effectively reducing overall run length.

### A purely static Tau obstacle, 3RS-Tau on subtilisin-treated microtubules, does not further impede KIF1A motility

In order to ascribe diffusive Tau as the inhibitory binding state on KIF1A pausing and subsequent motility, we needed to rule out static Tau as a potential inhibitory binding state. Previously conducted studies have revealed that removal of C-terminal tails removes the diffusive behavior of Tau, and shifts the microtubule bound population to a static binding state (Figure 5A) [25]. As our previous studies have characterized KIF1A’s motility and pausing behavior on subtilisin-treated microtubules [10], we next assessed the motility and pausing of KIF1A on 3RS-Tau coated, subtilisin-treated microtubules (purely static Tau obstacles) to compare these findings to the motility and pausing of KIF1A on 3RS-Tau coated, untreated microtubules (a mix of diffusive and static obstacles).

**Figure 5.**
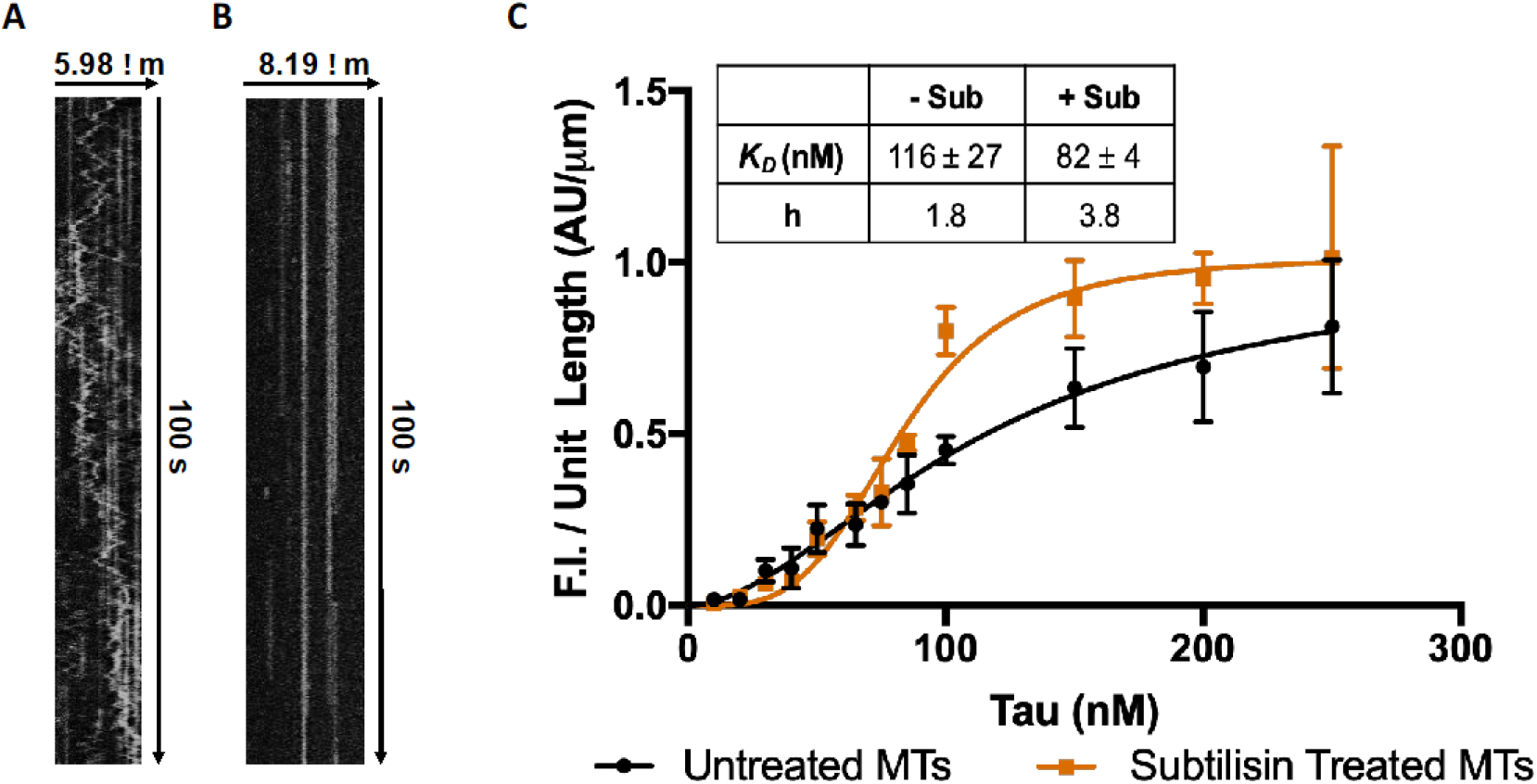
TIRF binding assays for Tau on untreated and subtilisin-treated microtubules (MTs). **A)** Representative kymograph of 3RS-Tau on untreated microtubules demonstrating the static (straight lines) and diffusive (jagged lines) behavioral states of Tau (160 pM Tau). **B)** Representative kymograph of 3RS-Tau on subtilisin-treated microtubules demonstrating that C-terminal tail removal shifts Tau to the static state (160 pM Tau). **C)** Upon subtilisin treatment of MTs, Tau’s affinity and cooperativity increased. 187 nM of 3RS-Tau was added to subtilisin-treated microtubules to account for differenced in binding affinity on untreated vs. subtilisin-treated microtubules, ensuring the same amount of 3RS-Tau decoration between these two microtubule populations.

First, to account for differences in 3RS-Tau binding affinity for the microtubule surface after microtubule C-terminal tail removal (Figure 5A-B), we performed a TIRF binding assay conducted as previously reported [16]. Upon removal of C-terminal tails, 3RS-Tau’s K_D_ decreased from 116 ± 40 nM to 70 ± 7 nM (Figure 5C). Additionally, we saw an increase in Hill coefficient from 1.8 to 4.1 (Figure 5C), suggesting that the removal of C-terminal tails results in an increase of 3RS-Tau’s cooperativity on paxlitaxel-stabilized microtubules. With this knowledge, we calculated the fraction of bound 3RS-Tau on subtilisin-treated microtubules at our highest experimental concentration of 200 nM. We then adjusted the concentration of 3RS-Tau added to ensure that the same fraction of 3RS-Tau is bound to subtilisin microtubules as our untreated microtubules, making shifts in 3RS-Tau’s behavioral state the only changing variable.

With the addition of 3RS-Tau to subtilisin treated microtubules (Figure 6A), we saw no significant change in KIF1A’s overall run length (3.26 ± 1.22 μm), overall speed (1.22 ± 0.23 μm/s), continuous run length (3.12 ± 1.12 μm), continuous speed (2.10 ± 0.47 μm/s), pause duration (1.34 ± 0.91 s), or pause frequency (0.17 pauses/overall run), when compared to KIF1A on subtilisin-treated microtubules without 3RS-Tau (Figure 6B-D, Figure S1 through S4, Table 1). Intriguingly, the addition of 3RS-Tau to subtilisin-treated microtubules significantly reduced the landing rate (1.92 ± 0.40 events/μm/min) of KIF1A in comparison to subtilisin-treated microtubules without Tau (Figure 4, Table 1), indicating that 3RS-Tau can reduce KIF1A landing rate independently of the C-terminal tail structure. Taken together, our results imply that the diffusive state of Tau relies on an interaction with the C-terminal tails to regulate KIF1A motility and behavior on the microtubule, and that the statically bound state of Tau does not regulate KIF1A motility or pausing.

**Figure 6.**
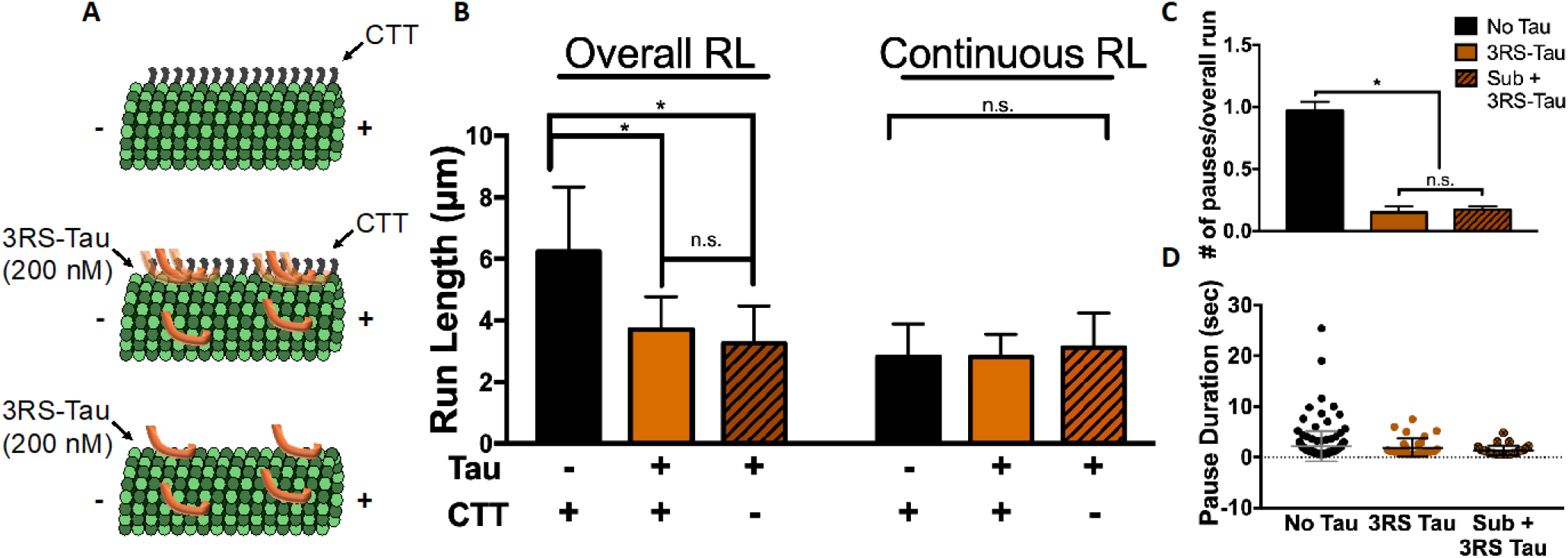
Addition of 3RS-Tau to subtilisin treated microtubules does not further inhibit KIF1A motility or pausing. **A)** Cartoon depiction of experimental conditions including microtubules without Tau and with C-terminal tails (CTT; top), microtubules with 3RS-Tau added (200 nM) and with CTTs (middle), or microtubules with with 3RS-Tau added (200 nM) and no CTTs (bottom). **B)** KIF1A overall run length was significantly reduced upon the addition of 3RS-Tau, or the addition of Tau on subtilisin (Sub)-treated microtubules, when compared to control microtubules (untreated, no Tau). KIF1A continuous run length was not significantly changed upon the addition of Tau, or the addition of Tau on subtilisin-treated microtubules, when compared to control microtubules (untreated, no Tau). **C)** KIF1A pause frequency (# of pauses/overall run) decreased on subtilisin-treated + 3RS-Tau microtubules (200 nM 3RS-Tau) when compared to control (no Tau, no subtilisin treatment) but not when compared to untreated microtubules with 3RS-Tau (200 nM 3RS-Tau). **D)** KIF1A pause duration on control microtubules, 3RS-Tau coated microtubules (200 nM 3RS-Tau) and subtilisin-treated + 3RS-Tau microtubules (200 nM 3RS-Tau). Characteristic run length values and standard deviations were calculated as previously reported [28].* p <0.05

## DISCUSSION

The significance of KIF1A’s role in long-range anterograde axonal transport [5, 11, 29, 30] is highlighted by the impact of altered KIF1A motility in the disease state [31-33]. Nevertheless, how this long-range transport is regulated for precise spatiotemporal delivery of cargo is poorly understood. The microtubule associated protein (MAP) Tau is an attractive possibility due to its disease state relevance and regulation of kinesin-1 motility [14-16, 18], yet the effects of Tau on KIF1A motility and behavior are relatively unexplored. Given our lab’s recent study detailing the necessity of tubulin’s C-terminal tails for KIF1A pausing behavior (Figure 7A) [10], we considered that Tau could regulate KIF1A motility in two mutually non-exclusive ways: 1) as an obstacle that truncates processive motility, and/or 2) by rendering the C-terminal tail inaccessible, prohibiting KIF1A from pausing. Here we present a new mechanism of Tau-mediated inhibition unique to KIF1A’s superprocessive motility and behavior. First, we reported the inhibitory capabilities of two Tau isoforms, 3RS-and 4RL-Tau, on KIF1A overall run length. In exploring this finding further, we then demonstrated Tau’s inhibitory effect occurs during characteristic KIF1A pausing behavior [10], supporting the idea that a reduction in overall run length results from a Tau-mediated reduction in number of pauses. Finally, we were able to ascribe this inhibition specifically to the diffusive binding state of Tau (Figure 7B).

**Figure 7.**
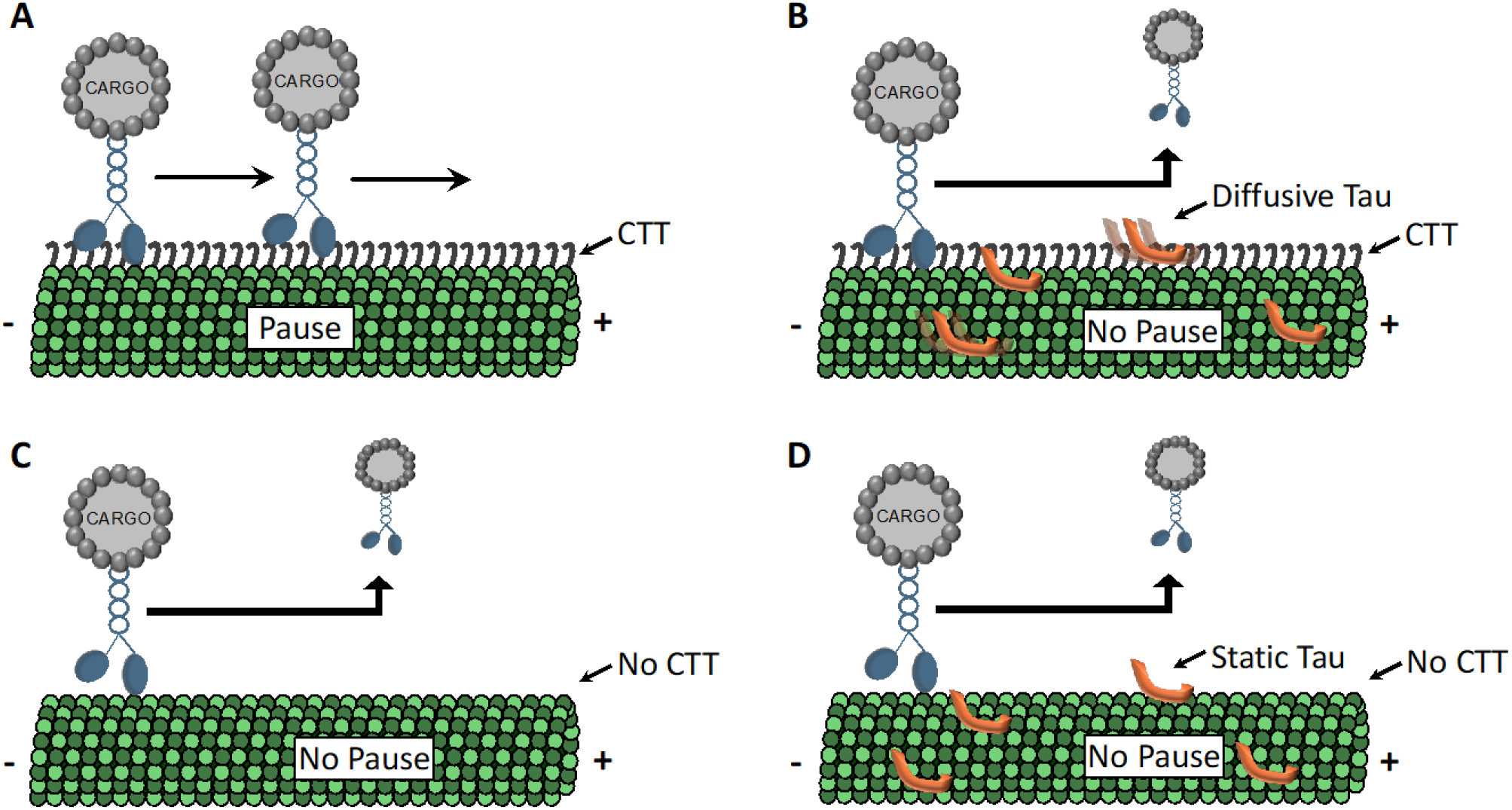
Model of Tau-mediated KIF1A regulation. **A)** In the absence of Tau, KIF1A exhibits an overall run length composed of continuous runs connected by pauses. These pauses are reliant upon accessible C-terminal tails (CTTs) to tether the KIF1A K-loop to the microtubule, before initiating another continuous run. **B)** When Tau is bound to the microtubule surface, the diffusive state of Tau interacts with and occupies the C-terminal tail. KIF1A can still achieve an initial continuous run, and navigate around static Tau obstacles, but cannot interact with the C-terminal tails due to diffusive Tau occupancy. This reduction in pausing decreases KIF1A’s ability to string together multiple continuous runs within an overall run length event. **C)** Tau’s inhibitory behavior on KIF1A is mimicked when C-terminal tails are proteolytically removed from the microtubule surface. A continuous run length can be initiated however, in the absence of C-terminal tails, the motor cannot tether to the microtubule to pause and is unable to string together multiple continuous runs within an overall run length event (as shown in [10]. **D)** Addition of Tau to subtilisin-treated microtubules results in a predominantly static population of Tau on the microtubule surface. Similar to panel B, KIF1A maintains the ability to navigate around static Tau obstacles, however, the absence of C-terminal tails limits the motors ability to pause. This reduction in pause frequency results in the inability to string multiple continuous runs together in an overall run length event.

When developing our model of Tau-mediated KIF1A regulation (Figure 7), it was important to consider how Tau is known to regulate other kinesin motor families. Foundational publications investigating Tau’s regulatory role on microtubule based motors revealed that Tau inhibits the processive motility of kinesin-1 motors by acting as an unnavigable obstacle, and that potency of inhibition varies by Tau isoform [14, 15]. Building upon these findings, our lab was able to ascribe Tau’s inhibition of kinesin-1 to Tau’s static binding state, not the diffusive binding state [16, 18]. In contrast, kinesin-2 motors are known to be insensitive to Tau, as their long neck linker allows them to switch microtubule protofilaments and side-step around obstacles [18, 19, 34].

If Tau regulates KIF1A as a static obstacle while the motor is processing (like kinesin-1 motors), we would expect to have seen a reduction in continuous run length upon addition of Tau, as this parameter is defined as segments where, within a single motility event, KIF1A is moving at a constant velocity [10]. However, our study revealed that, while the overall run length of KIF1A is reduced upon addition of either 3RS-or 4RL-Tau, the continuous run length does not change between experimental Tau concentrations (Figure 2). This same trend applies when looking at Tau’s effect on KIF1A overall vs continuous speed. As Tau concentration increased, and pause frequency was reduced, we observed an increase in overall speed (Table 1, Figure S1). This is likely resulting, in part, from a decrease in pausing events that would bring the overall speed down. In contrast, the continuous speed of KIF1A remains largely unchanged between 3RS-Tau concentrations. We did observe a decrease in continuous speed upon increase concentrations of the more diffusive 4RL-Tau isoform (Table 1, Figure S2). This leads us to postulate that diffusive Tau engaged with the CTT may not only inhibit KIF1A pausing, but may also interact with the KIF1A motor domain resulting in a slight impairment of speed.

Furthermore, when the C-terminal tails of tubulin were removed, biasing 3RS-Tau’s binding state to be exclusively static, there was no significant change in KIF1A motility or pausing behavior when compared to 3RS-Tau coated microtubules with CTTs still present (Figure 6, 7D). Thus, while this study revealed Tau’s ability to regulate KIF1A, it does not do so by acting as a static roadblock. Alternatively, these findings lead us to postulate that, like the kinesin-2 family, KIF1A is able to navigate around static Tau obstacles, likely resulting from differences in its neck linker structure [19, 35]. The concept of KIF1A using protofilament side-stepping to navigate around Tau is further supported by recent cryo-electron microscopy reconstructions of Tau on the microtubule surface, revealing that Tau binds along the crest of a single protofilament [36]. The assessment of KIF1A’s ability to side-step, and particularly how this behavior correlates with KIF1A’s neck linker length/sequence, is an enticing project for future studies in our lab.

Our current results support a model where Tau regulates KIF1A by rendering the C-terminal tail inaccessible, prohibiting KIF1A from pausing and connecting multiple continuous runs to generate a superprocessive overall run length. We postulate that this is the same mechanism responsible for inhibiting KIF1A landing rate, as both scenarios are mediated by KIF1A’s interaction with the tubulin C-terminal tails (CTTs). Past work in our lab has characterized KIF1A’s reliance on CTTs to engage in characteristic pausing behavior on the microtubule surface (Figure 7A, 7C) [10]. Furthermore, Tau relies on tubulin’s CTTs to engage in its diffusively bound behavioral state (Figure 5A-B) [25]. In further support of our model, both 3RS-and 4RL-Tau isoforms exhibited dosage dependent inhibition on KIF1A pause frequency (Figure 3, Figure S6), further confirming that KIF1A is not being regulated in the moments in which it is engaged in processive motility. Specifically, as the more diffusive isoform, 4RL-Tau, has a greater inhibitory potency on KIF1A pausing (Figure 3C-D) and overall run length when compared to the more static isoform, 3RS-Tau, at each experimental concentration (Figure 3C), we propose that the diffusive binding state of Tau is inhibiting KIF1A pausing behavior and subsequent motility. From our model, we can postulate that other MAPs known to engage with tubulin CTTs would have a similar effect on KIF1A function. Recent publications have begun to investigate this concept, revealing that not only is KIF1A sensitive to the presence of other MAPs, but the degree of inhibition is dependent on the combination of MAPs on the microtubule surface [20, 21].

Differences in affinity between Tau isoforms (Figure S7), or of Tau on different microtubule lattices, are imperative to consider when making dosage-dependent claims of Tau’s ability to regulate KIF1A motility. Consequently, a quantitative assessment of statically bound 3RS-Tau on subtilisin-treated microtubules (Figure 5B) was conducted over a range of concentrations (Figure 5C). These results demonstrated an increased Hill coefficient of 3RS-Tau on subtilisin-treated microtubules (Figure 5C) when compared to 3RS-Tau bound to untreated microtubules. While this increased cooperative behavior insinuates Tau’s ability to form patches, it is important to note that the lowest concentration our motility assays were conducted at is 50 nM, a concentration that is much higher than past literature where Tau patches were observed [14, 37]. Additionally, these experiments yielded an accurate *K*_*D*_ for 3RS-Tau on subtilisin treated microtubules (Figure 5C). Measuring Tau’s affinity in each of our experimental scenarios allowed us to calculate the fraction of Tau bound at given experimental concentrations, and adjust accordingly to ensure our KIF1A motors are encountering identical amounts of Tau bound to the microtubule surface.

Recent studies investigating Tau’s ability to form islands [38], patches [20], and condensates [37], have advanced our understanding of concentration dependent changes in Tau behavior and highlight an important structural state of Tau-mediated kinesin inhibition. As a result, it is important toconsider the dynamics of Tau molecules in the concentrations used in this study, and how these dynamics may differ in “low density” and “high density” environments. Previous work from our lab has assessed the isoform-specific Tau dynamics in low density and high density environments [26], revealing that, while the diffusion coefficient of individual Tau molecules decreases in high density environments, there is no change in dwell time. This means that an individual Tau molecule inside of a patch (or high density environment) would spend the same amount of time on the microtubule as an individual Tau molecule outside of a patch but would spend more time sampling the same area on the microtubule. In the context of this study, we propose that a patch would make the range of C-terminal tails less accessible for Tau engagement, but the dynamics of individual Tau molecules within the patch are unchanged.

With Tau further mechanistically characterized as a regulator of KIF1A motility, we can use our findings to further understand disease-state pathology. The regulatory capabilities of Tau’s static and diffusive binding behavior on axonal cargo transport are critical to understanding Tau’s role in neurodegeneration. In diseases known for impaired axonal transport, such as Alzheimer’s disease (AD) and frontotemporal dementia (FTD), changes in Tau isoform expression have been correlated to disease progression [22]. For example, pathological overexpression [39-41] of the more diffusive four-repeat Tau isoforms [26] and reduced transport of KIF1A cargo are observed in FTD [12, 13, 42]. Our results support a FTD disease state model in which a shift towards the diffusive state of Tau pathologically over-regulates KIF1A cargo transport in two ways. First, 4RL-Tau is significantly more inhibitory to KIF1A’s landing rate than 3RS-Tau (Figure 4), presenting an increased KIF1A “parking problem” [20]. Second, the KIF1A motors that are able to engage with the microtubule will have impaired motility due to the increase in diffusive Tau isoform expression, resulting in decreased pause frequency and overall run length. Additionally, the irregular localization and aggregation of KIF1A cargo seen in the neuronal pre-synaptic terminals in AD [11] can now be attributed to a loss of Tau-mediated KIF1A regulation, yielding “overactive” anterograde transport. This loss of regulation further explains an aggregation of KIF1A cargo at the distal neurite tips, as a reduction in microtubule bound Tau is observed in tauopathies resulting from hyperphosphorylation [23, 24, 43]. This logic is supported by recent work in *C. elegans* neurons, revealing that in PTL-1 (*C. elegans* Tau analog) knockdown worms, Unc-104 (KIF1A analog)-mediated synaptic vesicles travelled farther distances down the axon [44]. Additionally, a disease state loss of Tau on the microtubules can initiate signaling cascades known to alter the motility of *Drosophila* Unc-104 [45]. In summary, these findings bring clarity to the different pathological phenotypes of KIF1A dysfunction in AD and FTD, highlighting the physiological relevance of Tau’s ability to regulate cargo transport through isoform-specific behavioral equilibria and connecting Tau to KIF1A in neurodegenerative disease.

## ACKNOWLEGEMENTS

We thank David Warshaw and Guy Kennedy for training and use of the TIRF microscope at the University of Vermont. A special thank you to Vermont Livestock Slaughter & Processing (Ferrisburgh, VT) for supporting our work. This work was supported by National Institute of General Medicine Sciences/National Institutes of Health funding to C.B. (Grant GM132646).

## AUTHOR CONTRIBUTIONS

Conceptualization, investigation, methodology, draft composition and editing by D.V.L. and C.L.B. Data curation and formal analysis by D.V.L. Supervision, funding, and project administration by C.L.B.

