## Supplemental Methods and Data for "The Microtubule Associated Protein Tau Regulates KIF1A Pausing Behavior and Motility"

**Running Title:** Tau regulates KIF1A behavior & motility

**Authors:** DV Lessard, CL Berger.

**Materials Included:**
Supplemental Figures and Legends 1 through 7
Supporting References

**SI FIGURES**

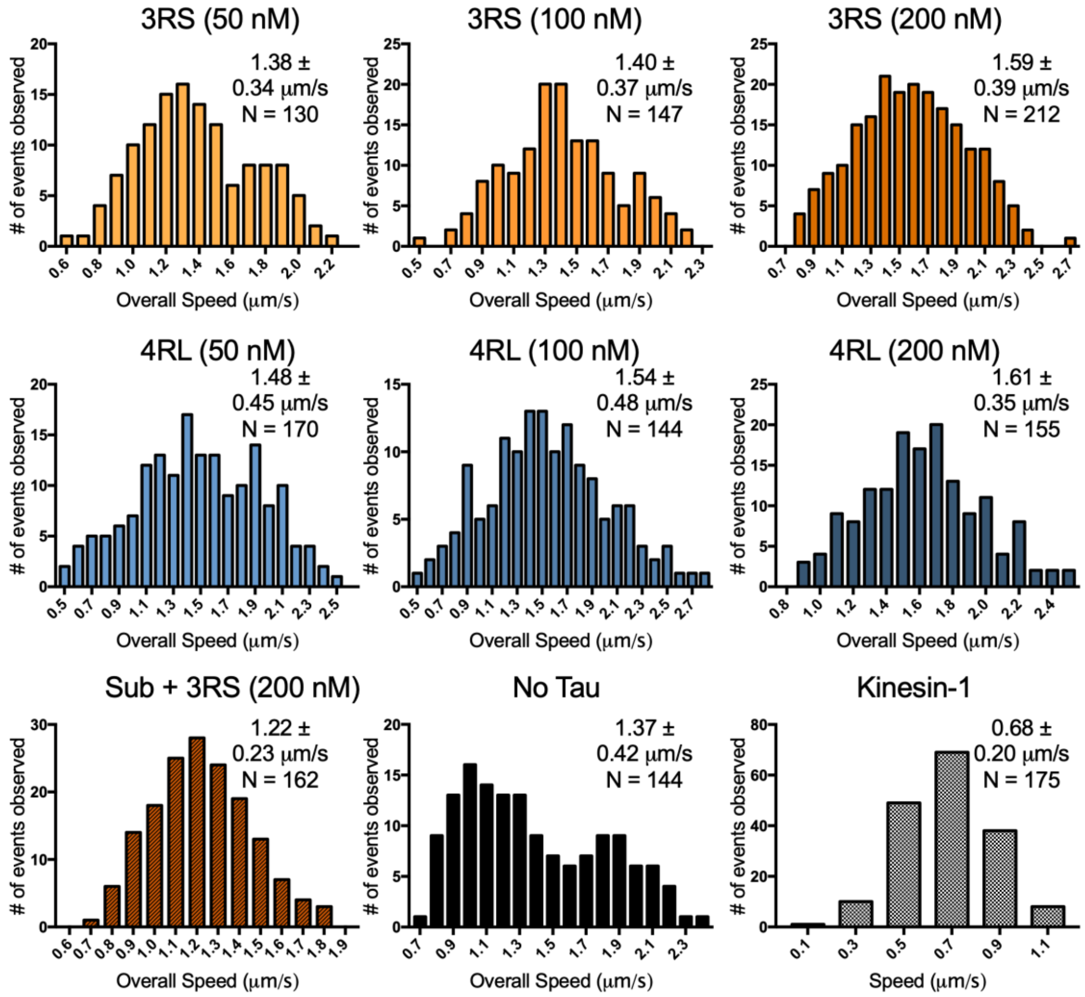


**Figure S1. Histograms of overall speed across all conditions (Mean ± SD).**

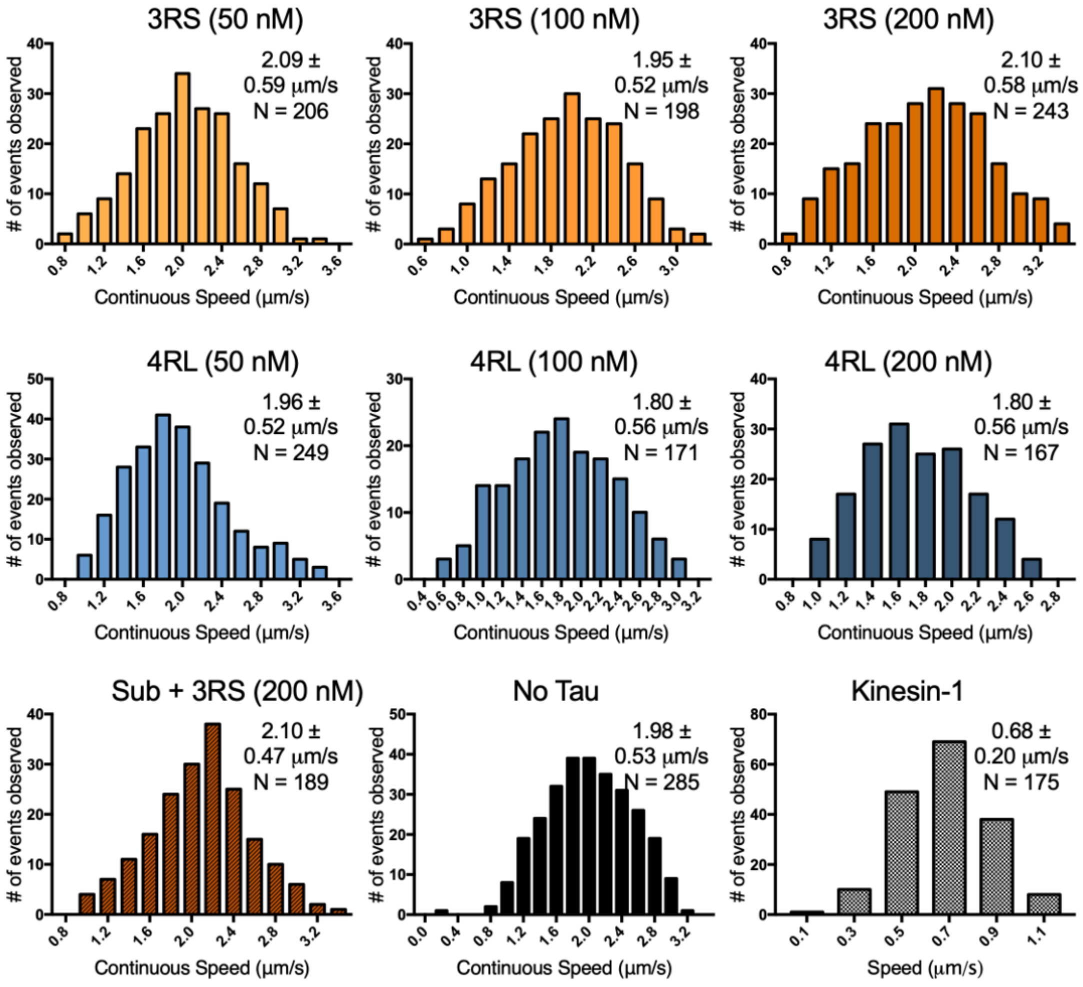


**Figure S2. Histograms of continuous speed across all conditions (Mean ± SD).**

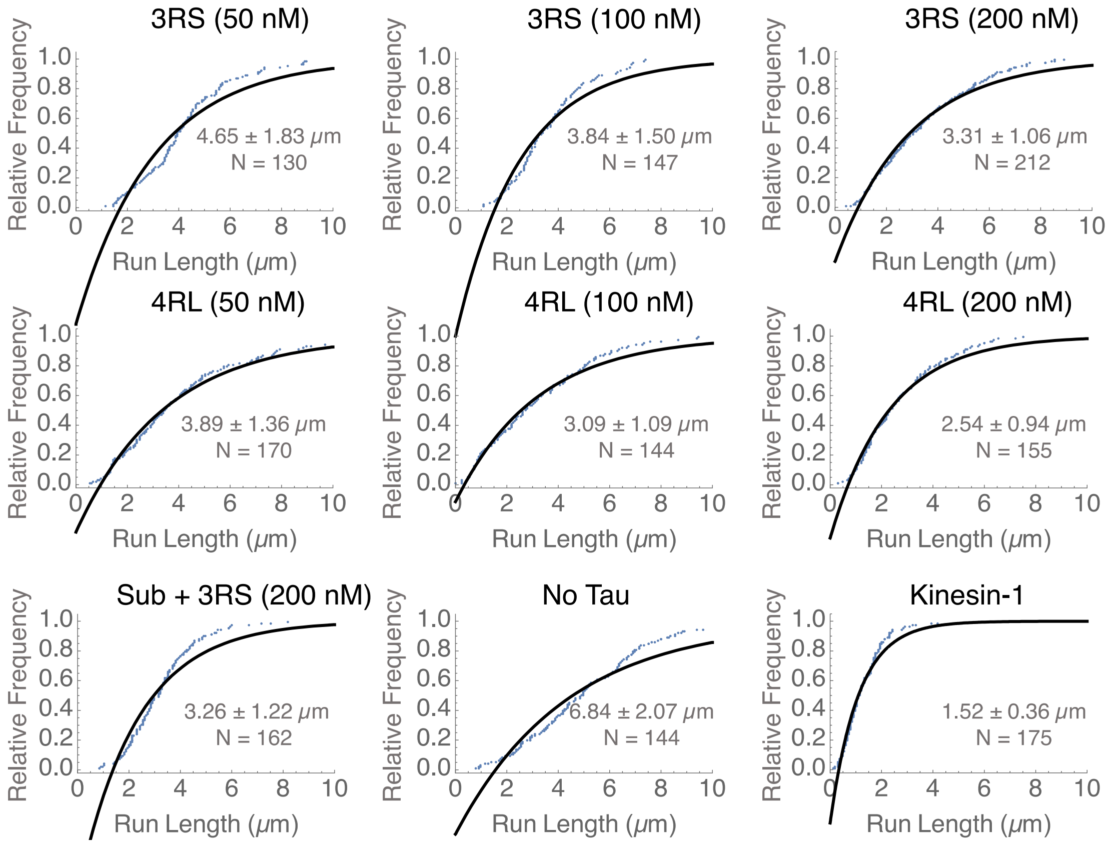


**Figure S3. Cumulative frequency plots representing the overall run length across all experimental conditions.** The raw run length is represented by black dots the observed cumulative frequency is represented by grey dots. Shown within each graph is the overall run length, calculated as previously reported [1] (Mean ± SD).

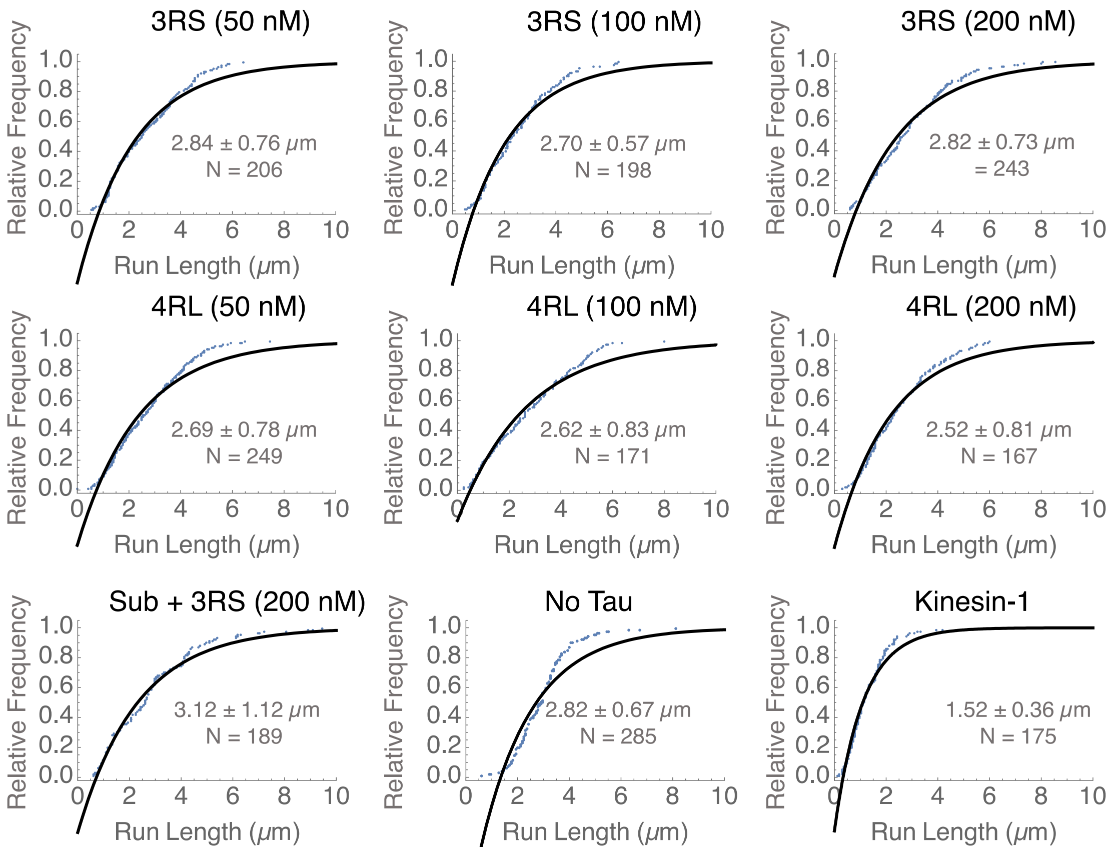


**Figure S4. Cumulative frequency plots representing the continuous run length across all experimental conditions.** The raw run length is represented by black dots the observed cumulative frequency is represented by grey dots. Shown within each graph is the overall run length, calculated as previously reported [1] (Mean ± SD).

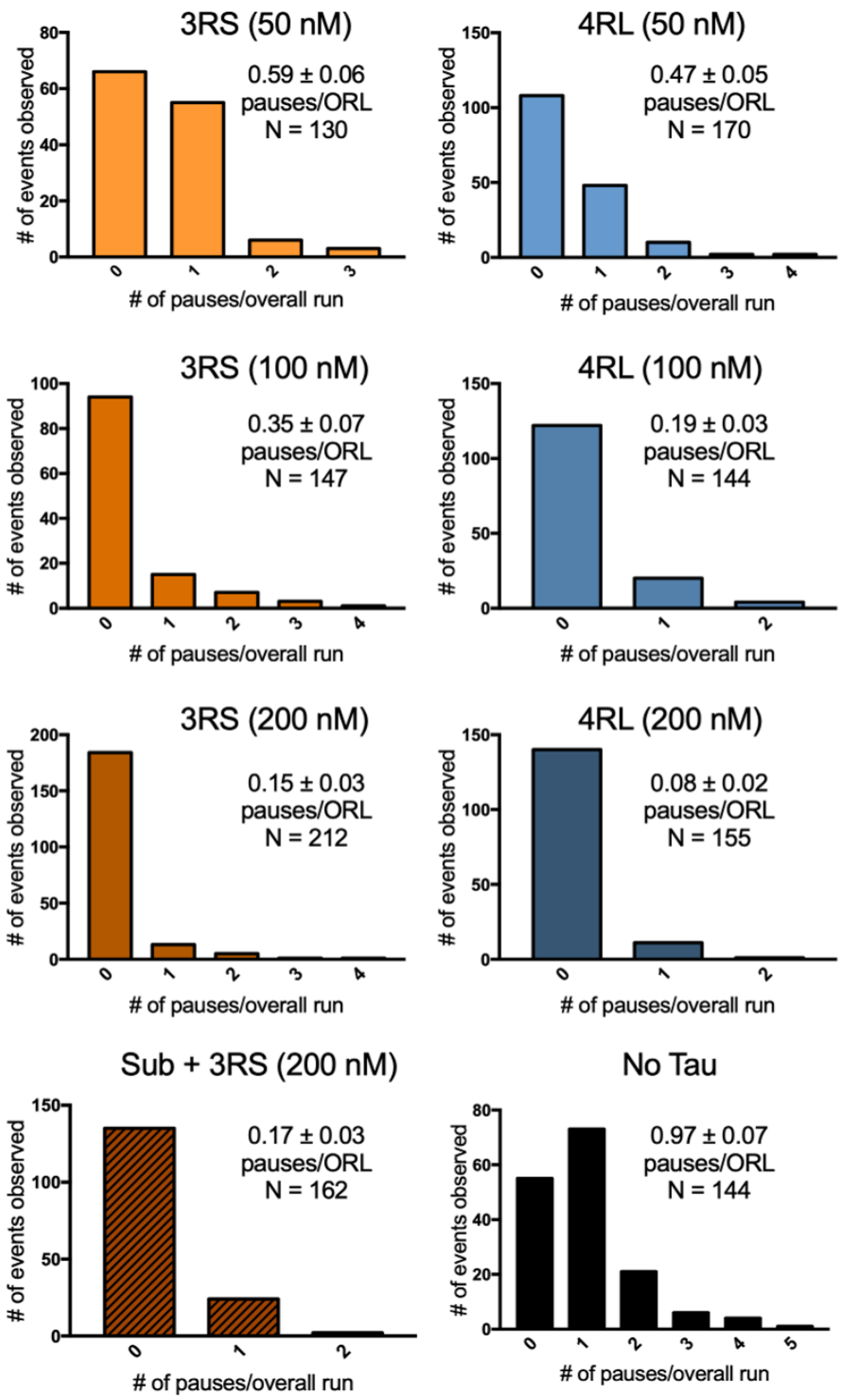


**Figure S5. Histograms of pause frequency (pauses/overall run [ORL]) across all conditions (Mean ± SEM).**

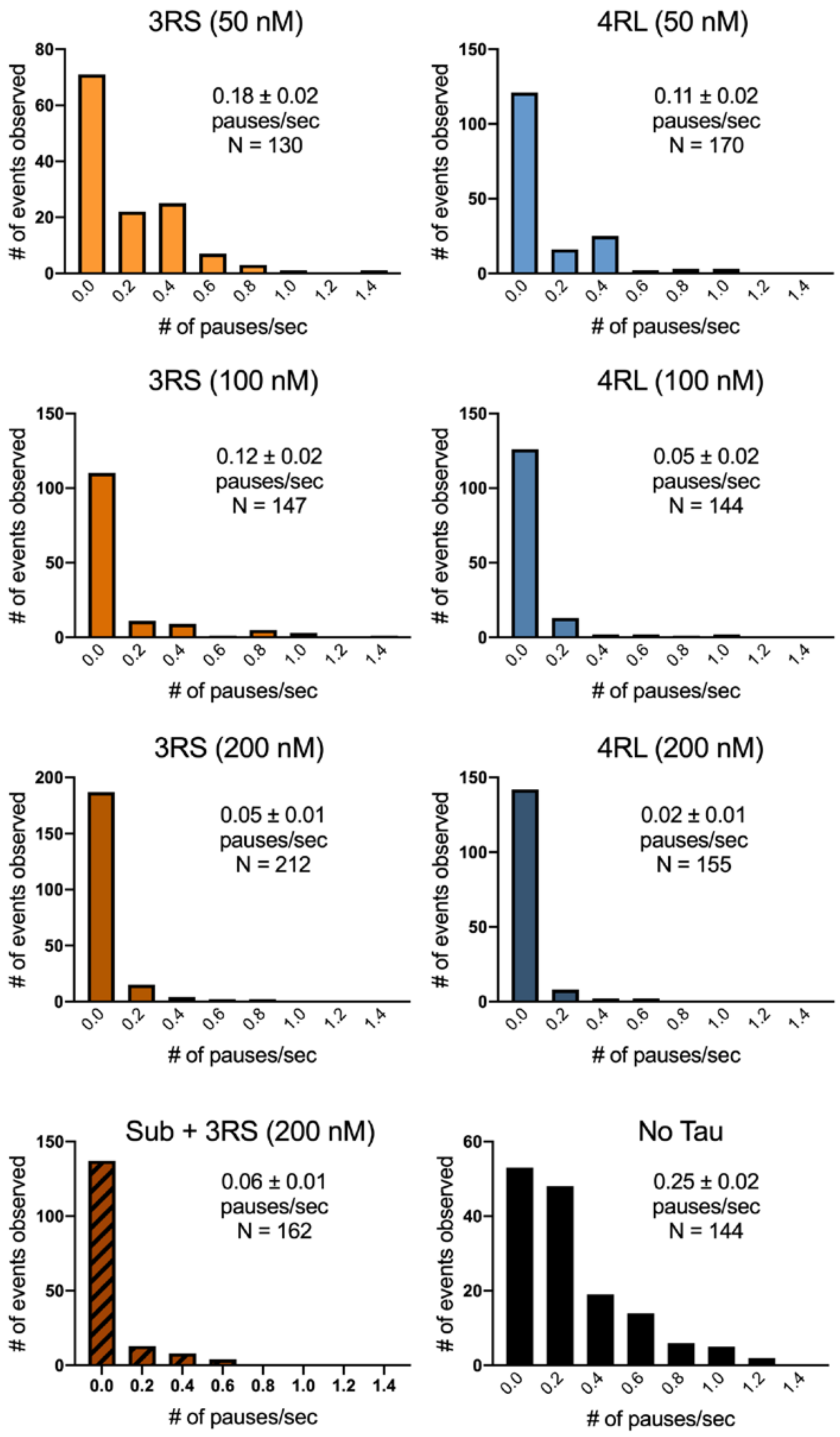


**Figure S6. Histograms of pausing per unit time (pauses/second) across all conditions (Mean ± SEM).**

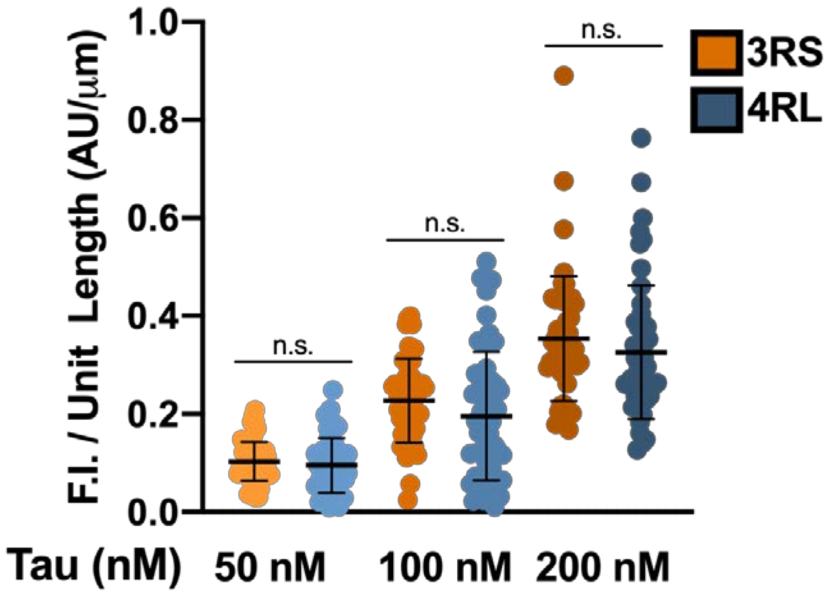


**Figure S7. Average fluorescence intensity per unit length between 3RS- and 4RL-Tau-coated microtubules at each experimental concentration (Mean ± SD).**
